# Ultra-long-range interactions between active regulatory elements

**DOI:** 10.1101/2022.11.30.518557

**Authors:** Elias T. Friman, Ilya M. Flyamer, Shelagh Boyle, Wendy A. Bickmore

## Abstract

Contacts between enhancers and promoters are thought to relate to their ability to activate transcription. Investigating mechanisms that drive such chromatin interactions is therefore important for understanding gene regulation. Here, we have determined contact frequencies between millions of pairs of cis-regulatory elements from chromosome conformation capture datasets and analysed a collection of hundreds of DNA-binding factors for binding at regions of enriched contacts. This analysis revealed enriched contacts at sites bound by many factors associated with active transcription. We show that active regulatory elements, independent of cohesin and polycomb, interact with each other across distances of 10s of megabases in vertebrate and invertebrate genomes and that interactions correlate and change with activity. However, these ultra-long-range interactions are not dependent on RNA polymerase II transcription or several transcription cofactors. We propose that long-range interactions between cis-regulatory elements are driven by three distinct mechanisms – cohesin-mediated loop extrusion, polycomb contacts, and association between active regions.

## Introduction

The large genomes of most animal species derive their regulatory potential from non-coding cis-regulatory elements (CREs), in particular the activation of genes by enhancers. The spatial relationship between CREs and transcription is a topic of active debate (see for example (Benabdallah et al., 2019; Karr et al., 2022; Mir et al., 2019; Xiao et al., 2021; Zuin et al., 2022)) and thinking in this area has been strongly influenced by genome-wide chromosome conformation capture methods such as Hi-C (Lieberman-Aiden et al., 2009) and micro-C (Hsieh et al., 2015). Contact frequency, which is the measure of how often two regions are in proximity for crosslinking and ligation, strongly scales negatively with the genomic distance between two regions (Lieberman-Aiden et al., 2009).

CREs controlling developmentally regulated genes are typically contained within a single topologically associating domain (TAD), formed by the highly dynamic process of cohesin-mediated loop extrusion (Gabriele et al., 2022; Symmons et al., 2014). Regions within a TAD interact more frequently with each other than with regions at similar distances outside the TAD. This is because cohesin can extrude chromatin until it reaches barrier elements, most notably CTCF, which delineate TAD boundaries (Fudenberg et al., 2016; Nora et al., 2017). Furthermore, focal contacts and “stripes” are seen between pairs of CTCF sites in the presence of loop extrusion (Rao et al., 2017). Both TADs and cohesin-driven focal contacts are limited to regions separated by up to 1-2 megabases (Mb) (Boyle et al., 2020; Dixon et al., 2012). Much less understood are mechanisms that lead to enriched interactions between regions over larger genomic distances, and which may be independent of cohesin.

One process bringing distal regions together is compartment interactions with their own chromatin type, within and between chromosomes. Initial analysis revealed A and B domains, corresponding roughly to active and inactive chromatin, that are several hundred kilobases (kb) to a few Mb in size (Lieberman-Aiden et al., 2009). More fine-scale mapping revealed “sub-compartments” within these larger domains (Rao et al., 2014; Spracklin et al., 2021), with extremely high sequencing depths revealing ‘compartments’ of sizes down to a few kb (Gu et al., 2021). A unified mechanism that explains compartment interactions has not been described, although perturbation of transcription in *Drosophila melanogaster* or DNA methylation in human cell lines can affect interactions between active and inactive compartments, respectively (Rowley et al., 2017; Spracklin et al., 2021). Compartment interactions may be driven at least in part by affinity between molecules differentially present in the compartments, such as histone marks, and may involve phase separation (Nichols and Corces, 2021; Nuebler et al., 2018).

In repressed chromatin, enriched focal interactions are detected between polycomb-bound regions over a very wide range of genomic distances, up to several 10s of Mb (Boyle et al., 2020; Loubiere et al., 2020; Rhodes et al., 2020). These interactions are cohesin-independent but are dependent on components of the polycomb repressive complexes. The functional significance of these focal interactions in polycomb-mediated repression is unclear (Dimitrova et al., 2022; Ogiyama et al., 2018).

There is also evidence for associations between active genomic regions over very large genomic distances. TADs containing super-enhancers and active genes are spatially closer to other highly active TADs than to those with low activity (Beagrie et al., 2017), and active regions tend to colocalise with nuclear speckles enriched in splicing factors (Chen et al., 2018; Quinodoz et al., 2018). In mouse embryonic stem cells (mESCs) and differentiated cell types, active transcription start sites (TSSs) and transcription factor (TF) binding sites have been shown to interact with their own type across 2-10 Mb (Bonev et al., 2017). The function of, and mechanism driving, these long-range interactions, and whether they also occur in other cell types and species, are not known.

To uncover mechanisms underlying enriched chromatin interactions between CREs at different scales of genomic separation, we performed a computational screen utilising micro-C and Hi-C data and ChIP-seq databases of binding sites for hundreds of DNA-binding factors in mouse and human cells. We searched for factors whose binding is correlated with enriched chromatin interactions between pairs of CREs at short (<1 Mb) and long (>1 Mb) genomic distances. In addition to polycomb and cohesin-associated proteins, we uncovered many proteins associated with active transcription whose binding sites were enriched in long-range interactions. These interactions at active genes and enhancers are enriched between regions separated by several 10s of megabases and even on different chromosomes, are dynamic between tissues and cell types, and occur in both vertebrates and invertebrates. We show that these interactions are unrelated to cohesin and polycomb-driven interactions, but also do not depend on transcription per se or several factors associated with transcriptional activation. We propose that CRE-CRE interactions are driven by three main mechanisms – loop extrusion, polycomb contacts, and ultra-long-range interactions between active regulatory elements.

## Results

### Screening for factors bound at sites of enriched CRE-CRE interactions

To quantify interactions between CREs, we merged DNase-accessible ENCODE (Luo et al., 2020) CREs from mESCs within 5 kb of each other and generated observed over expected contact frequencies for each CRE pair from Hi-C and micro-C data at both short-range (0.1-1 Mb) and long-range (1-10 Mb) (Figure 1A). The 10 Mb cut-off is to limit the number of combinations and to avoid sparse data at larger distances. We determined the overlap of CREs with DNA-binding sites (peaks) in the cistromeDB (Mei et al., 2017) and ReMap2022 (Hammal et al., 2022) databases. For each peak dataset representing one ChIP-seq experiment, we divided CREs into overlapping and non-overlapping. We then compared the contact frequencies of CRE pairs overlapping the factor on both sides or not at all by calculating the Mann-Whitney adjusted p-value and effect size (F=U/(n_1_*n_2_)), where F>0.5 means increased contact frequencies at bound compared to unbound (Figure 1B). We annotated the factors into 7 classes; cohesin-associated, polycomb-associated, transcription cofactors, TFs, repressive factors, and other. We use transcription cofactors broadly here to mean chromatin and transcription-associated proteins which are not TFs or known repressors.

**Figure 1.**
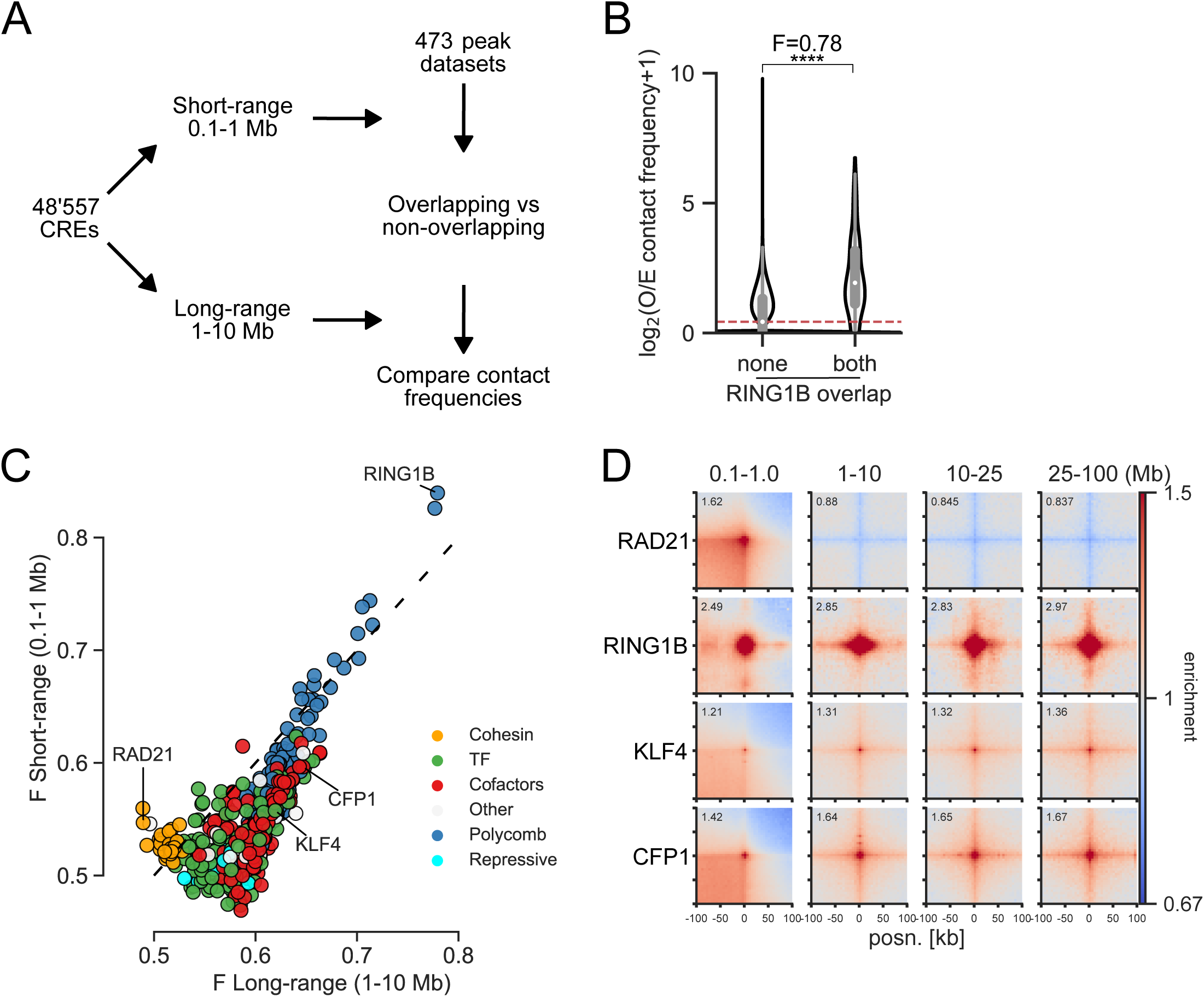
Computational screening for factors associated with enriched interactions between CREs. **(A)** Schematic of the computational pipeline. CRE pairs are split into short-range (0.1-1 Mb between pairs) and long-range (1-10 Mb between pairs). These regions are overlapped with peaks from different datasets and contact frequencies at 10 kb resolution compared between pairs overlapped by the factor, and pairs not overlapped by the factor to calculate enrichment values. **(B)** Example of the analysis for a RING1B ChIP-seq dataset. Observed over expected (O/E) contact frequencies for CREs at long-range were split into overlapping RING1B peaks on neither or both sides and Mann-Whitney F and adjusted p-values were calculated (****: p=3.8*10^-184^). **(C)** Effect sizes for factors with significantly enriched chromatin interactions compared to unbound CREs in mESCs. x and y axes show enrichment at short-range and long-range. Colours represent the group the factor belongs to. **(D)** Pileup analysis for 5000 regions for each of the indicated hits from Fig. 1C for interaction pairs at different distances of genomic separation in micro-C data from mESCs. For CFP1 and KLF4, peaks overlapping RING1B binding sites were excluded.

We tested our approach by measuring changes in enrichment in the seven different classes in micro-C and Hi-C datasets from cells in which cohesin-associated factors had been degraded (Hsieh et al., 2021) or Ring1b (polycomb) knocked out (Boyle et al., 2020). As expected, RAD21 and CTCF degradation led to a decrease (Supp. Fig. 1A,B), and WAPL degradation led to an increase (Supp. Fig. 1C) in enrichment between cohesin-associated binding sites at short-range. Ring1b knockout (KO) led to an expected decrease in enrichment for polycomb-bound regions at both short and long range (Supp. Fig. 1D).

After confirming that our approach was able to detect factors associated with enriched chromatin interactions, we performed our analysis on micro-C data from WT mESCs (Hsieh et al., 2020) (Fig. 1C, Supp. Fig. 2A-B, Supp. Table 1). Cohesin-associated factors were enriched for interactions at short but not long range, and polycomb-associated factors were similarly enriched at short and long range. Besides these already known mechanisms, binding sites for transcription cofactors and TFs tended to be enriched at both short and long range, but with higher enrichment at long range. We correlated the number of non-polycomb bound TSSs per CRE bound by each factor and its enrichment. TFs and cofactors had a high correlation between enrichment and TSS overlap, showing that they are separate from polycomb and related to the presence of genes (Supp. Fig. 2C). We also performed the same enrichment analysis on Hi-C data from the GM12878 human lymphoblastoid cell line (Rao et al., 2014) and saw a similar pattern of enrichment for the different classes, meaning that this is not a feature specific to micro-C data or to mESCs (Supp. Fig. 2D-F). We used pileup analysis (Flyamer et al., 2020) to confirm some of the top hits. Sites bound by the CpG island (CGI) binding protein CFP1 (CXXC1), or KLF4 - a pluripotency TF active in mESCs, showed enrichment at all distances up to 100 Mb (Fig. 1D). We call this non-cohesin, non-polycomb associated category of interactions at large distances “ultra-long-range interactions” (ULIs) between active regions.

ULIs are seen as enriched stripes and central pixels in pileups, meaning that these active elements interact with each other and with the surrounding chromatin more than other regions at similar distances. Note that while the level of enrichment above expected is similar across distances, this does not represent similar absolute contact frequencies. We considered potential technical artifacts that could lead to the appearance of ULIs. We divided accessible regions into quartiles based on DNase-seq signal, split them into those within 1kb of a TSSs or at least 5kb from the nearest TSS, and generated pileups (Supp. Fig. 3A-B). While ULIs scaled with accessibility, TSSs in lowly accessible regions were much more enriched in ULIs than more accessible sites without TSSs, arguing against this signal simply being a result of increased digestion or crosslinking efficiency in accessible chromatin. We also considered that large protein complexes bound at active genes could lead to increased crosslinking efficiency. However, contact frequencies did not correlate with the number of cistromeDB and ReMap2022 peaks overlapping CREs, suggesting this is not the case (Supp. Fig. 3C-D). We also excluded normalisation artifacts, as ULIs can be seen in both unbalanced and balanced (iterative convergence and eigenvector decomposition, ICE, normalised, (Imakaev et al., 2012)) data, whether balanced on all, or only on cis, contacts (Supp. Fig. 3E). ULIs are also seen regardless of normalising by expected values, shifted controls, or not at all. We therefore conclude that ULIs represent bona fide enriched contacts between active regulatory elements.

### Ultra-long-range interactions between active regions are independent of cohesin and polycomb

The strong enrichment for CFP1 binding sites in our screen (Fig. 1C, Supp. Fig. 2B) prompted us to test the relationship between CGIs and ULIs. This showed a correlation between interactions and the density of CpG dinucleotides at CGIs devoid of polycomb (Fig. 2A). Interaction strength at TSSs also scaled strongly with the level of transcription (Fig. 2B). Notably, TSSs in quartile 4 (highly transcribed) showed enrichment with quartiles 3 and 2 but not with quartile 1, meaning that inactive genes do not interact at all (Supp. Fig. 4A). Non-CGI promoters also interact but much less than CGI promoters, suggesting CGIs are not required but contribute to ULIs at TSSs (Supp. Fig. 4B). Analysis of micro-C data at 100 bp resolution within the 10kb region surrounding the TSS showed that enriched interactions are centred at the TSS (Fig. 2C).

**Figure 2.**
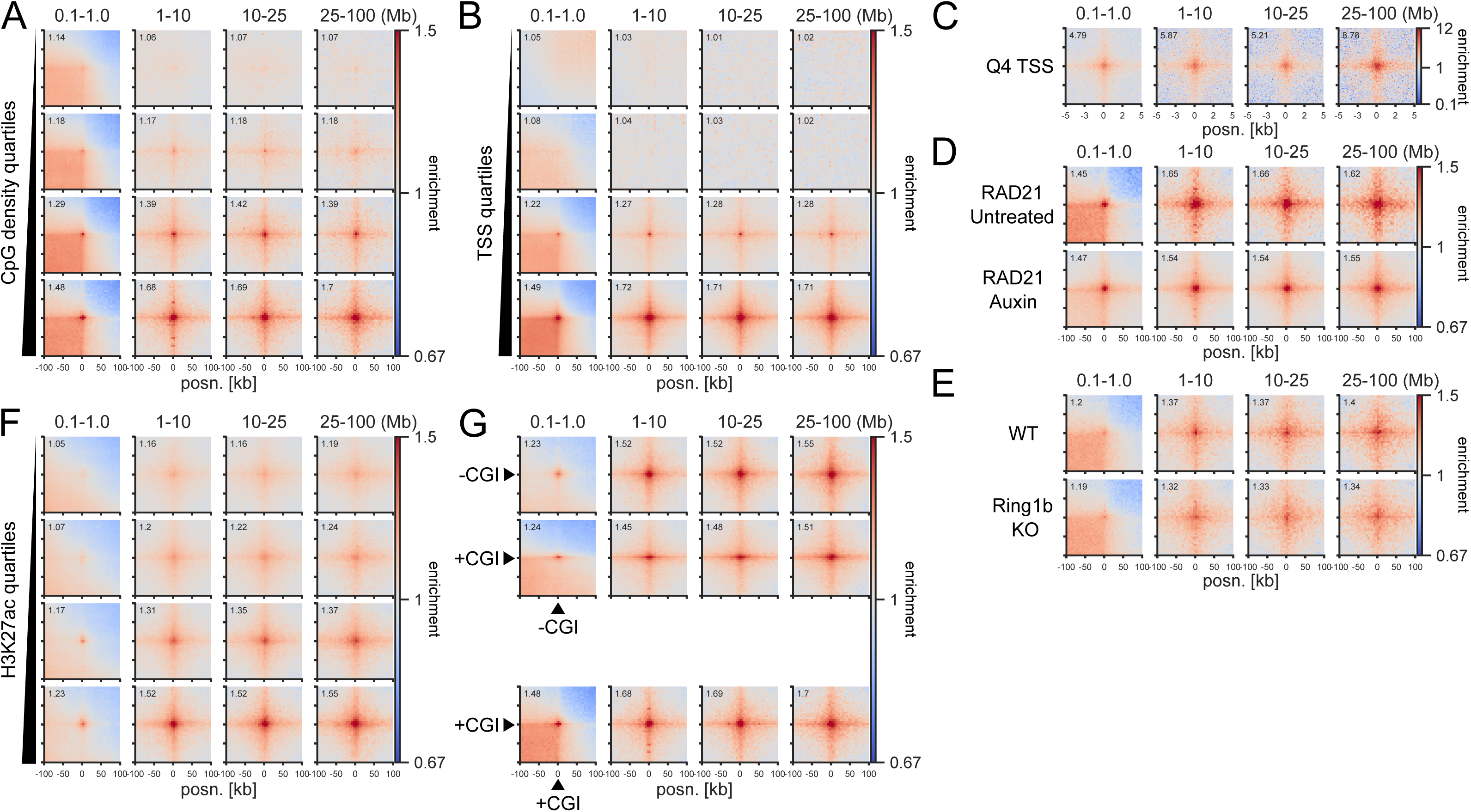
Ultra-long-range interactions between active promoters and distal regulatory elements. Pileup analysis for **(A)** non-RING1B overlapping CGIs split by quartiles based on CpG density (2352-3015 peaks per group). **(B)** Non-RING1B overlapping TSSs (4800-5128 peaks) divided into quartiles by expression level, based on 4SU-seq data, in micro-C data from mESCs. **(C)** The 10 kb surrounding the Q4 TSSs (5128 peaks) at 100 bp resolution in micro-C data from mESCs. **(D and E)** CGI Q4 (2651 peaks) regions in **(D)** micro-C data from RAD21-AID mESCs and **(E)** Hi-C data from WT or Ring1b KO mESCs. **(F)** TSS-distal H3K27ac peaks not overlapping RING1B or CGIs split by quartiles based on H3K27ac ChIP-seq signal (6161-6452 peaks per group) in micro-C data from mESCs. **(G)** CGI Q4 regions (2651 peaks, +CGI) and Q4 promoter-distal H3K27ac (6268 peaks, -CGI) in in micro-C data from mESCs.

To formally exclude that ULIs are dependent on polycomb and cohesin, we analysed data from mESCs with Ring1b deleted (KO) or RAD21 degraded (AID) for 3 hours, which disrupts polycomb interactions and CTCF-CTCF interactions, respectively (Boyle et al., 2020; Hsieh et al., 2021). These interventions did not affect ULIs (Fig. 2D-E), and neither did 3 hours of depletion of CTCF or the cohesin unloader WAPL (Supp. Fig. 4C). However, we did note a strong reduction of interactions after 6 hours of RAD21 depletion in an independent mESC dataset (Rhodes et al., 2020) (Supp. Fig. 4D), but not in data from Rad21 KO mouse thymocytes (Seitan et al., 2013) (Supp. Fig. 4E).

We reasoned that this could be related to cell cycle dynamics as, unlike thymocytes, mESCs divide rapidly and accumulate in G2/M after 6 hours of RAD21 depletion due to failure to proceed through mitosis (Rhodes et al., 2020). Analysis of a Hi-C dataset from cell cycle-synchronised erythroblasts (Zhang et al., 2019) showed, as expected, that ULIs are lost in mitosis and reappear as early as anaphase/telophase (Supp. Fig. 5A). This is prior to the formation of both CTCF loops and compartments (Zhang et al., 2019). Thus, the loss of ULIs after 6 hours of RAD21 depletion is most likely explained by the accumulation of mitotic cells in the absence of cohesin function.

Analysis of cell-cycle phased merged single-cell Hi-C data (Nagano et al., 2017) shows that ULIs persist throughout G1, S, and G2 phases (Supp. Fig. 5B). In human HAP1 cells (Haarhuis et al., 2017), we did not detect a loss of ULIs upon deletion of the cohesin subunit SCC4, while WAPL deletion led to slightly decreased interactions (Supp. Fig. 5C). It is possible that the stiffening of the chromatin fibre that occurs upon prolonged loss of WAPL (Tedeschi et al., 2013) leads to a less dynamic chromatin structure, preventing active regions from interacting.

Promoter-distal H3K27ac-marked CREs, i.e putative enhancers, which do not overlap with CGIs, also show enriched ULIs that scale with the level of H3K27ac enrichment (Fig. 2F). ULIs also occurred between these distal H3K27ac regions and CGIs as well as expressed genes (Fig. 2G, Supp. Fig. 5D). Taken together, these results show that ULIs scale with the activity of regulatory elements, are independent of polycomb and cohesin, are present in cycling and non-cycling cells, are reformed quickly after mitosis, and persist throughout the cell cycle.

### Rare interactions between distal cis-regulatory elements

Enrichment in the pileups represents averages of many pairs of regions. High average enrichment could come from a few highly interacting pairs or from a tendency of many or all pairs to interact. To test this, we assessed the distribution of contact frequency values using two approaches, focusing on TSS quartile pairs at 10-25 Mb distance. First, we quantified the contact frequency in the central 5 kb bin containing the TSS. Higher expression quartiles had higher contact frequencies, although most values were zero in all quartiles, reflecting the sparsity of data at these large distances (Fig. 3A). We also looked at the signal of the individual “corner stripes” in the pileups (Fig. 3B). This revealed a higher number of non-zero values across the stripe with increased expression level (Q3 and Q4). We did not find evidence of a few particularly highly interacting pairs dominating the pileups, showing rather that this is a property of increased interaction frequencies between many pairs.

**Figure 3.**
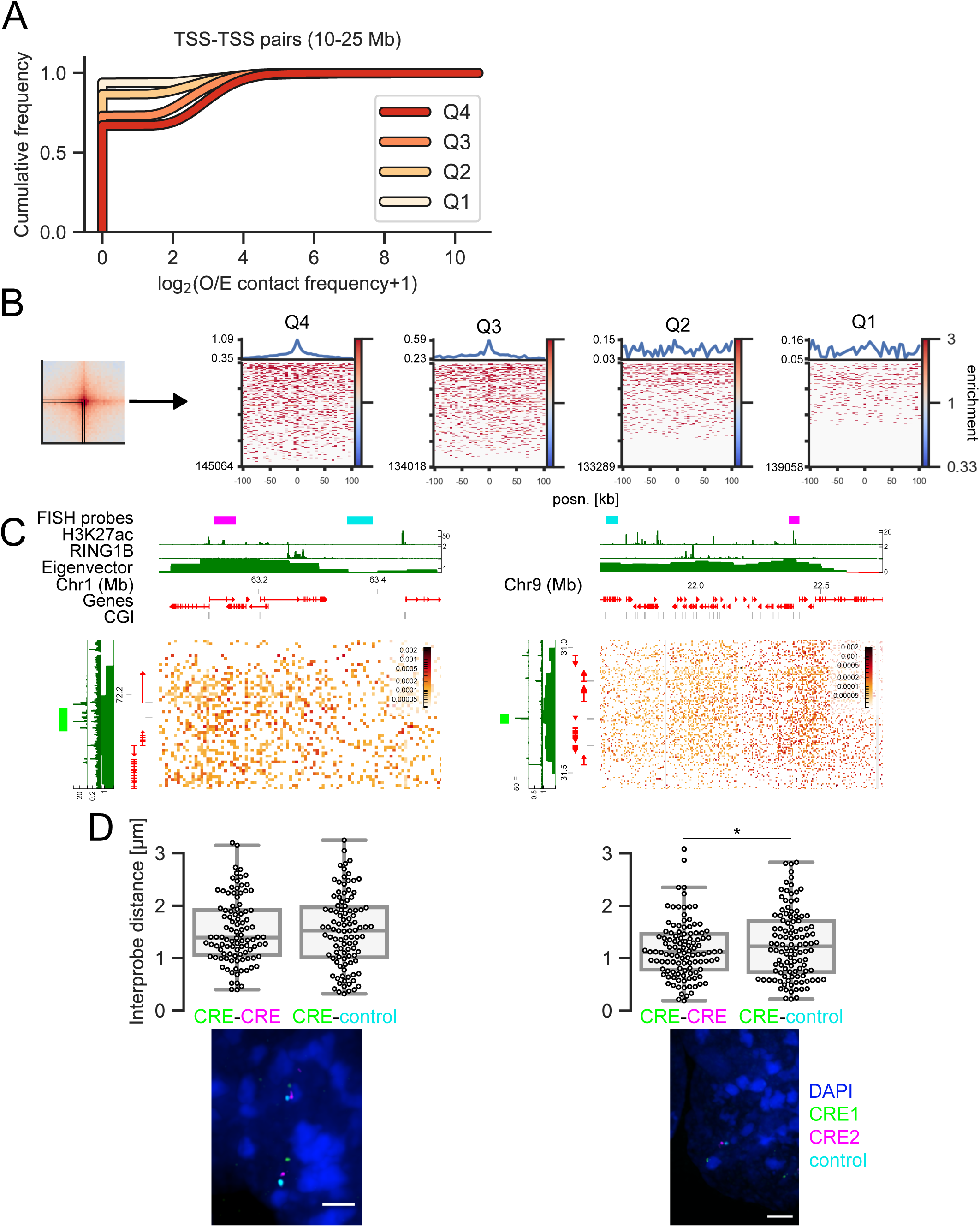
Distribution of contact frequencies and DNA FISH. **(A)** Empirical cumulative distribution frequency of O/E contact frequency values in the central 5 kb pixel between TSSs in the different quartiles separated by 10-25 Mb. **(B)** Left: Depiction of the “corner stripe” from a pileup. Right: Heatmaps of O/E contact frequency values in individual corner stripes for TSS-TSS quartile interaction pairs at 10-25 Mb distance. The line plot on top represents the mean signal. **(C)** Hi-C matrices for regions on chromosomes 1 (left) and 9 (right) used in DNA FISH experiments. CGI marks CGIs without RING1B. **(D)** Top: Interprobe distances measured by DNA FISH between ∼9 Mb distal CREs on chromosomes 1 (left) and 9 (right) or a CRE and an equidistant control region in the same A compartment. N=100 (Chr1) and 112 (Chr9) measurements. Boxplot lines denote maximum (excluding outliers), interquartile range (upper and lower bound of box), median (centre of box), and minimum values. *: p=0.03. Bottom: Example images of DNA FISH. Blue: DAPI, Green: CRE1 probe, Magenta: CRE2 probe, Cyan: control probe (adjacent to CRE2 in the genome). Scale bar: 2 μm.

The analyses above show that ULIs represent higher average contact frequencies between active CREs compared to the surrounding chromatin. Contact frequencies are related to, but are not a direct measure of, spatial distances (Finn et al., 2019). Nevertheless, we expected that active CREs separated from each other by large genomic distances would be closer together in the nucleus on average than with regions at similar distances that are not active CREs. To test this, we performed DNA FISH on mESC nuclei using probes between pairs of CREs at 9 Mb of genomic separation or between one CRE and an equidistant control region in the same A compartment. Both CRE-CRE pairs had lower median distances separating them, than the corresponding CRE-control, though for only one of these was this difference signficant (p=0.03, significant at FDR=10% after Benjamini-Hochberg correction) (Fig. 3C). This indicates that while the average enrichment between distal CREs is high in pileups, this only translates to marginally smaller distances between individual distal CREs in nuclei (median distances in both cases was > 1 micron). We only detected one instance of colocalisation (<200 nm distance) between 212 measured CRE-CRE distances, showing that the enriched contacts seen between many regions in Hi-C/micro-C are exceedingly rare between individual pairs (see Discussion).

### Ultra-long-range interactions occur primarily in *cis* and in the A compartment

Because ULIs are enriched at such large genomic distances, we tested if they are also enriched in *trans* between chromosomes. Indeed, high CpG-density regions devoid of polycomb are enriched in *trans* interactions (Supp. Fig. 6A). We note that while the level of enrichment compared to expected is comparable to that within chromosomes, the absolute contact frequencies are very low. We also considered that interactions might occur between homologs and tested this using Hi-C data from hybrid mESCs with phased SNPs (Han et al., 2020). This showed that interactions occur primarily within chromosomes and not between homologs (Supp. Fig. 6B).

Active regions in the A compartment at distances of several 10s of Mb, as those we look at here, will be separated by multiple intervening A and B compartments. We wondered if the stripe seen in the pileups would span the whole intervening chromatin or be constrained to A compartments. We picked regions close to A/B compartment switches and saw that the interaction enrichment was confined to the A compartment, i.e. the stripe does not span into the B compartment (Supp. Fig. 6C).

### Interaction dynamics and DNA methylation

Their relationship to transcriptional activity and H3K27ac suggests that ULIs are dynamic between tissues. To investigate this, we used data from mESCs and differentiated neural progenitor cells (NPCs) (Bonev et al., 2017) and selected H3K27ac peaks enriched in either of the cell states (Fig. 4A). These regions showed correspondingly higher enrichment of ULIs in the cell state with higher H3K27ac. To investigate more rapid dynamics, we examined Hi-C data from human macrophages stimulated with lipopolysaccharide (LPS) and interferon (IFN-γ) (Fig. 4B) (Reed et al., 2022). Regions that gained H3K27ac upon stimulation also gained ULIs and regions losing H3K27ac, although relatively lowly enriched to begin with, lost ULIs over the time course. While the small number of regions affected and the time resolution of the experiment precludes delineating which changes come first, it appears that changes in H3K27ac and ULIs accompany one another.

**Figure 4.**
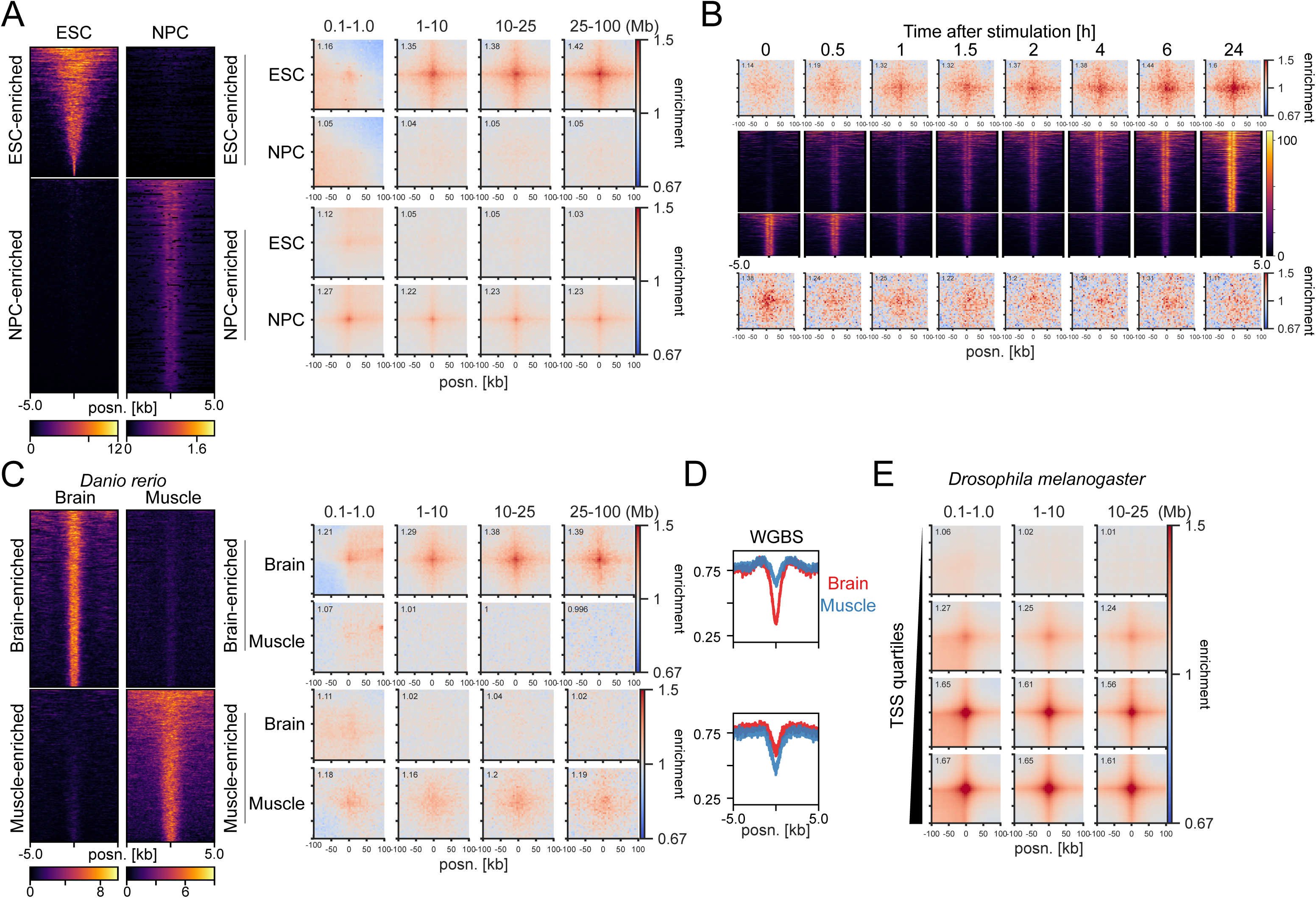
Interaction dynamics in mouse, human, zebrafish and *Drosophila melanogaster*. **(A)** Left: H3K27ac signal in mESCs and neural progenitor cells (NPC) at mESC-enriched (1509 peaks) and NPC-enriched (2491 peaks) regions. Right: Pileup analysis for the differentially enriched H3K27ac peaks in Hi-C from mESC and NPC. **(B)** Top and bottom: Pileup analysis for regions with increasing (top; 1058 peaks) or decreasing (bottom; 553 peaks) H3K27ac across the time course of LPS and IFN-γ stimulation of THP-1 derived macrophages. Middle: H3K27ac signal in THP1-derived macrophages at region with increasing (top) or decreasing (bottom) H3K27ac. **(C)** Left: H3K27ac signal in zebrafish brain and muscle at brain-enriched (2315 peaks) and muscle-enriched (1994 peaks) regions. Middle: Pileup analysis for the differentially enriched H3K27ac peaks in brain and muscle. **(D)** Average whole genome bisulfite (WGBS) signal in the differentially enriched H3K27ac peaks in brain and muscle. **(E)** Pileup analysis for *Drosophila melanogaster* TSSs (2943 regions per group) divided by expression level based on RNA-seq in Hi-C data from *Drosophila melanogaster* eye-antennal imaginal discs.

To examine dynamic changes in ULIs between tissues *in vivo*, we took advantage of data from *Danio rerio* (zebrafish) and selected brain- or muscle-enriched promoter-distal H3K27ac peaks (Yang et al., 2020). These were accompanied by enrichment of ULIs in the respective tissue (Fig. 4C). We also saw a corresponding higher level of DNA methylation in the tissue where the regulatory elements were inactive and no interactions were seen (Fig.4D). Because of the relationship between CGIs and ULIs (Fig. 2A), we reasoned that the focal demethylation at these regulatory elements may be responsible for the interactions, and that gain in DNA methylation would lead to their loss. To test if methylation is required for ULIs, we used Hi-C data from *Drosophila melanogaster*, a species with little DNA methylation and lacking CGIs (Deaton and Bird, 2011). We could detect ULIs between expressed TSSs and H3K27ac-positive, non-polycomb bound (as well as polycomb bound), regions in *D. melanogaster* eye-antennal imaginal discs (Loubiere et al., 2020) (Fig. 4E and Supp. Fig. 6D). This shows that ULIs are not specific to vertebrates and that DNA methylation is not required for their formation.

### Ultra-long-range interactions are not directly dependent on transcription

In our initial screening, we had seen enrichment for the binding sites of many sequence-specific TFs at ULIs (Fig. 1C, Supp. Fig. 2A-B). To determine whether homotypic interactions between TFs could drive ULIs we performed enrichment analysis comparing enrichment values for sites where a particular TF is bound at both interacting sides to where that TF is bound at only one side but there is at least one other TF bound at the other side. This is to avoid comparing to regions devoid of binding altogether at one side, i.e. inactive regions, which would skew the enrichment. Binding at both sides yielded a stronger enrichment than at one side, but the two were correlated with significant enrichment for every TF also when bound at only one side (Fig. 5A). As an example, we split H3K27ac peaks into those bound by MYC or not in the GM12878 lymphoblastoid cell line and saw that MYC-bound regions interacted with other active regions not bound by MYC (Supp. Fig. 7A). Although we cannot exclude some contribution, this analysis indicates that homotypic TF-TF interactions are unlikely to drive ULIs.

**Figure 5.**
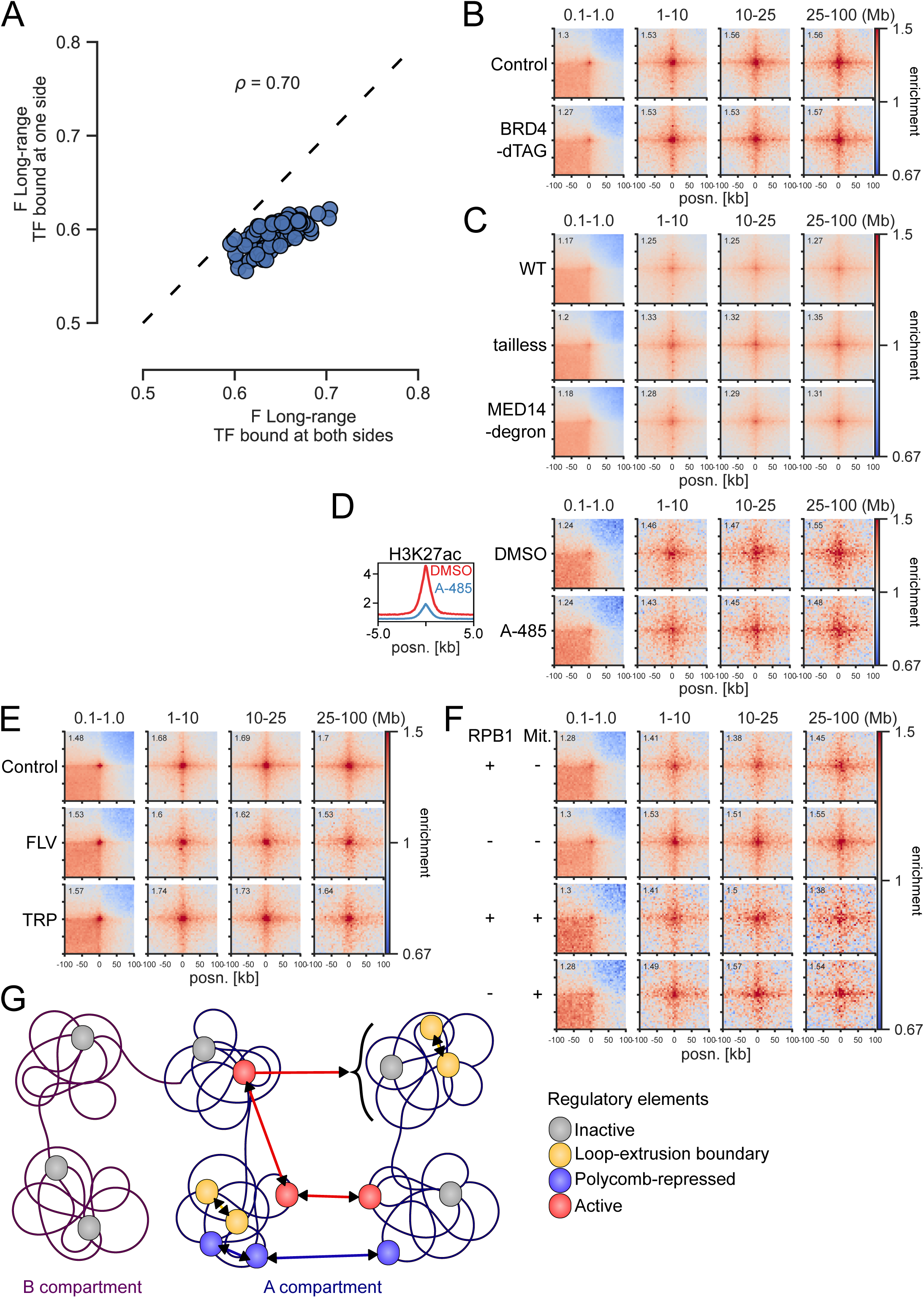
Independence of BRD4, Mediator, P300 activity, and transcription. **(A)** Effect sizes of TFs for long-range (1-10 Mb) contact enrichment compared to TF-unbound CREs in mESCs. x and y axes show enrichment for TF binding at both sides and only one side, but at least one other TF bound at the other side. ρ: Spearman’s correlation coefficient. **(B)** Pileup analysis between CGI Q4 regions in dTAG-BRD4 mESCs with or without BRD4. **(C)** Pileup analysis between CGI Q4 regions in WT, Med15^-/-^ Med16^-/-^ Med23^-/-^ Med24^-/-^ Med25^-/-^ (tailless), and MED14-SMASh degron CH12 cells. **(D)** Left: Mean H3K27ac signal in CGI Q4 regions in mESCs treated with DMSO or A-485. Right: Pileup analysis between CGI Q4 regions in mESCs treated with DMSO or A-485. **(E).** Pileup analysis between CGI Q4 regions in micro-C data from WT mESCs or treated with flavopiridol (FLV) or triptolide (TRP) for 45 minutes. **(F)** Pileup analysis between CGI Q4 regions in RPB1-AID mESCs with or without RPB1. Top two rows represent asynchronous cells, bottom two rows represent cells arrested in mitosis and released into G1. **(G)** Proposed model for interactions between CREs. Inactive genes in A or B compartments do not interact. In the A compartment, CREs overlapping loop-extrusion boundaries interact via cohesin at short range, and polycomb-repressed CREs interact at short and long range. Active regulatory elements interact with each other across very large distances (focal interactions), as well as with other A compartments (stripes).

We next tested if transcription cofactors may be required for ULIs. BRD4 has been implicated in chromatin organisation (Linares-Saldana et al., 2021; Rosencrance et al., 2020). Furthermore, Mediator and BRD4 have both been shown to be unevenly distributed in the nucleus, and enriched in so-called “transcriptional condensates” (Cho et al., 2018; Sabari et al., 2018). However, our analysis shows that either BRD4 degradation (Linares-Saldana et al., 2021) or Mediator disruption (El Khattabi et al., 2019) do not affect ULIs (Fig. 5B,C). YY1 has been implicated as a regulator of enhancer-promoter contacts (Weintraub et al., 2017), but degradation of YY1 (Hsieh et al., 2021) also had no effect on ULIs (Supp. Fig. 7B). Treatment with the P300 catalytic inhibitor A-485 (Pelham-Webb et al., 2021) leads to loss of P300 activity and H3K27ac, but has no effect on ULIs (Fig. 5D).

We considered whether ULIs may be driven by association with nuclear speckles, as this is strongly correlated with gene activity (Chen et al., 2018). Knockdown of the splicing component Srrm2 leads to a partial disruption of speckles (S. Hu et al., 2019), but this did not grossly affect ULIs (Supp. Fig. 7C). Nuclear speckle association is primarily seen in the A1 subcompartment (Chen et al., 2018). We split active regions (H3K27ac peaks) in GM12878 by A1 and A2 (Rao et al., 2014) association and saw a higher level of ULI enrichment in A2 (Supp. Fig. 7D). These data suggest that speckles cannot fully explain ULIs, although we cannot exclude that they may have some contribution.

Active regulatory elements, including CGIs, tend to be preferentially located toward the nuclear interior (Beck et al., 2018; Boyle et al., 2001). To test whether ULIs are a reflection of a more central nuclear position of these regions, we used GPSeq data from HAP1 cells which measures radial positioning genome-wide (Girelli et al., 2020). We divided GPSeq regions into three bins (1-3), from the periphery to the centre of the nucleus. As expected, CGIs lacking H3K27me3 in HAP1 cells were enriched in the most central GPSeq bin (Supp. Fig. 7E). Analysis of pileups between regions in the three bins revealed no trend towards more interactions between centrally occupying regions, in *cis* or in *trans* (Supp. Fig. 7F). However, this analysis normalises the signal to the corners of the pileup to see enrichment compared to surrounding chromatin. When excluding this step and looking simply at average observed over expected contact frequencies in the different bins, interactions are highest between regions in the most central bin at the largest genomic distances (25-100 Mb) and in *trans* (Supp. Fig. 7G). These higher contact frequencies surrounding the CGIs likely reflects a higher level of A compartment intermingling. Our analysis indicates that (i) interactions between CGIs happen more often in the centre of the nucleus because they tend to be localised there, (ii) focal enrichment in contact frequencies between CGIs over the surrounding chromatin is similar regardless of nuclear radial positioning measured by GPSeq, and (iii) *trans* or very distal *cis* interactions are higher between regions in more centrally occupying regions. We therefore conclude that radial positioning contributes to, but does not fully explain, ULIs.

As levels of transcription are correlated to ULIs, we investigated whether loss of transcription would disrupt ULIs. Treatment with Flavopiridol or Triptolide, which block transcription elongation and initiation, respectively, for 45 minutes did not lead to a loss of interactions (Hsieh et al., 2020) (Fig. 5E). Neither did degradation of RPB1, the largest subunit of RNA polymerase II (Pol II), for 6 hours (Jiang et al., 2020) (Fig. 5F). ULIs are lost in mitosis and regained before cells enter G1 (Supp Fig. 5A). When analysing cells that were synchronised in mitosis and then released into G1 with or without RPB1, we saw no loss of ULIs (Fig. 5F).

## Discussion

Overall, our results indicate that although ULIs involving active CREs interacting across large genomic distances are characterised by scaling with activity – levels of transcription or H3K27 acetylation – they are not dependent on RNA Pol II or several other cofactors involved in transcriptional regulation, including BRD4, Mediator, and P300 activity.

ULIs are dynamic, changing between cell types and upon stimulation on the timescale of a few hours. There must therefore be some driving event leading to ULIs, which also coincides with activation of the genomic loci involved. We cannot exclude that there is a single factor mediating these interactions which we have not uncovered. However, we find it more likely that ULIs reflect an emergent property that is related to the many events that are involved in driving CRE activity, such that disrupting any individual factor does not disrupt the tendency of active regions to interact with each other. The fact that interactions are re-established after mitosis without a functional Pol II shows that ULIs are not merely a consequence of ongoing transcription.

Many other studies have observed CRE-CRE interactions at distances above 1 Mb (for example (Bonev et al., 2017; Ogiyama et al., 2018; Schoenfelder et al., 2015)). Those long-range interactions which cannot be explained by polycomb likely reflect the same phenomenon we describe here, i.e. the tendency of active regions to interact. CGI-containing genes have been shown to interact more than non-CGI genes within and across chromosomes (Beck et al., 2018). This was attributed to their more central nuclear localisation and interaction with either transcription factories or polycomb bodies. We do not find a dependency of active ULIs on transcription components, nor a direct relationship to radial positioning. A caveat of this analysis is that the measure of radial positioning we used (GPSeq) represents an average of the entire cell population and has relatively low resolution (100 kb). Interactions between promoter CGIs and non-promoter (orphan) CGIs have been implicated in driving transcriptional activation, although at much shorter genomic distances (Pachano et al., 2021). While ULIs are particularly strong at CGIs, we see interactions between CGIs and other non-CGI active regions. While our results do not exclude some level of specificity between different classes of active regions, we did not see evidence of this.

Two recent studies describe small-scale “compartments” uncovered at very high sequencing depth using either whole-genome Hi-C, or micro-C with capture of individual TADs (Goel et al., 2022; Gu et al., 2021). While we propose ultra-long-range as a defining feature of active CRE-CRE interactions, we do not exclude that the same forces driving them operates also at shorter distances (i.e. within TADs). Indeed, our initial screen showed enrichment for factors associated with active regions both at short-range and long-range, with a correlation between the two. While the compartment nomenclature used in the above studies suggests that these interactions are formed by the same mechanism as the much larger compartment domains, it is not yet known exactly what mechanisms contribute to compartment interactions. We observe that ULIs reform after mitosis before G1, which is much earlier than the reestablishment of large A/B compartments, which are complete only by late G1. While this does not exclude a similar mechanism, it suggests that ULIs behave differently from the larger compartment domains.

DNA FISH of CRE pairs 9 Mb apart showed only one instance of colocalisation (<200 nm) of spots and marginally smaller median distances compared to CRE-control probes in the same A compartment. Therefore, ULIs should not be considered as stable interactions between individual regions, but rather an increase of already small probabilities of association between CREs compared to surrounding chromatin. This could derive from an increased probability for encounter in the nucleus, an increase in time spent together when encountered, or both. It is the cumulative increase in these probabilities between many regions which give rise to the average enrichment in Hi-C/micro-C data.

We propose the following model for interactions between CREs across scales (Fig. 5G). At the ∼1-2 Mb scale CREs are brought into proximity by the loop extrusion process, with focal interactions between loop extrusion boundaries such as CTCF sites. Polycomb-repressed regions are both locally compacted and interact with each other across all genomic distances. Finally, active CREs, including promoters and enhancers, interact with each other across large genomic distances. The average enrichment between active CREs is on the order of ∼50-100%, which could be considered relatively low. It is hard to imagine that very rare contacts between two specific regions at large distances, even though higher than with other regions at similar distances, would have a measurable functional impact. However, the cumulative interactions between all active CREs are substantial and may have a functional effect by creating a nuclear environment that enhances transcription.

## Methods

### Custom code

All custom code used in the analysis has been deposited on GitHub (https://github.com/efriman/Friman_etal_ULI). Custom analysis was done in Python 3.8.12 (G. van Rossum (Guido), 1995) in Jupyter notebooks (Kluyver et al., 2016) using pandas (McKinney, 2010), numpy (Harris et al., 2020), and scipy (Virtanen et al., 2020) for data analysis and matplotlib (Hunter, 2007) and seaborn (Waskom, 2021) for plotting.

### Hi-C data processing

Already processed Hi-C and micro-C data were, when required, converted to cooler format (Abdennur and Mirny, 2020) using hic2cool (Dekker et al., 2017) or cooler cload pairs (see Supp. Table 3). Hi-C pairs from allele-phased SNPs (Han et al., 2020) were downloaded from NCBI GEO:GSE132898 and split into separate files by genotype annotation, followed by conversion to cooler using cooler cload pairs. Other datasets (see Supp. Table 4) were processed using the distiller-nf 0.3.3 (Goloborodko et al., 2019) pipeline with default settings and coolers filtered for q>=30 were used. ICE balancing (Imakaev et al., 2012) was performed using cooler balance, using ‘--cis_only’ or ‘--trans-only’ to generate cis/trans balanced weights. Replicate datasets were combined when available using cooler merge. Expected contact frequencies were generated using the cooltools 0.5.1 (Open 2C et al., 2022) functions expected-cis or expected-trans with ‘--clr_weight_name’ set to the appropriate balancing (cis/trans/total). Hi-C data in Fig. 3C were visualised using HiGlass in resgen.io (Kerpedjiev et al., 2018).

### Contact screen

ReMap2022 peaks were downloaded from https://remap2022.univ-amu.fr/. cistromeDB “Factor” data for mouse and human were downloaded in batch from http://cistrome.org/db. cistromeDB datasets corresponding to perturbation datasets (e.g. siRNA, KO, treatments) or misannotated data were discarded (see Supp. Table 5). ReMap2022 and cistromeDB data for mouse or human were combined and overlapped with ENCODE CREs (peaks within 5kb merged using BEDTools merge (Quinlan and Hall, 2010)) overlapping DNase-seq peaks in mESCs or GM12878, respectively (see Supp. Table 6). Observed over expected (O/E) contact frequencies at 10 kb resolution (balanced using all contacts) and their respective coordinates for DNase-accessible CREs were extracted using coolpup.py with settings ‘-- store_stripes --flank 0 --expected expected_file --mindist 100000 --maxdist 10000000 --by_distance 100000 1000000 10000000’. For each dataset in the cistromeDB and ReMap2022 data the contact frequency scores for short-range and long-range were split between pairs overlapping the dataset at both ends or at neither end. Datasets with less than 500 regions or with less than 50 overlapping or non-overlapping regions were discarded. A Mann-Whitney U test (using scipy.stats.mannwhitneyu with default settings) was performed to compare the two distributions and the effect size (F) calculated using F=U/n_1_*n_2_ where n_1_ and n_2_ are the number of observations for overlapping and non-overlapping regions. Adjusted p-values were calculated by multiplying the Mann-Whitney p-value by the number of tests performed. TFs and transcription cofactors were derived from the AnimalTFDB 3.0 (H. Hu et al., 2019) database followed by additional manual annotation (see Supp. Table 1-2). For Supp. Fig. 1, the F values were normalised to the mean F in each dataset (e.g. Ring1B KO) and this normalised value used to divide treated over untreated or KO over WT. For Fig. 5A, the analysis was performed in the same way as described above, except peaks were combined for all ChIP-seq datasets of the same factor. Comparison was made between regions unbound by any TF to those bound on both sides by the factor or only one side but at least one other TF bound on the other side.

### Pileups

Pileups were generated with coolpup.py 1.0.0 (https://github.com/open2c/coolpuppy) (Flyamer et al., 2020). Most pileups are generated with the command ‘coolpup.py coolfile peakfile --flank 100000 --mindist 100000 --maxdist 100000000 --by_distance 100000 1000000 10000000 25000000 100000000 –expected expected_file.tsv’. Balancing (ICE normalisation) was always done on cis contacts, except for Supp. Fig. 6A where trans contacts were used, and Supp. Fig. 6B where all contacts were used. For all pileups, 5 kb resolution data were used, except for Fig. 2C where 100 bp resolution was used. For Supp. Fig. 3E, mindist was set to 1000000 and -- clr_weight_name was altered based on if using total, cis, or no balancing, and ‘-- nshifts 0’ ‘--nshifts 5’ or ‘--expected expected_file’ used. Central pixel interactions and stripes in Fig. 3A-B were generated using ‘--mindist 10000000 --maxdist 25000000 --store_stripes’. For Supp. Fig. 6C, regions near compartment edges were annotated as + or - stranded depending on if the nearest boundary was to the left or right and pileups generated with ‘--by_strand --mindist 1000000 --maxdist 100000000 --flip_negative_strand’ and combining -- with ++ using a custom modify_2Dintervals_func in coolpup.py. For figures comparing interactions between different sets of regions, e.g. Fig. 2G, the different sets were annotated as + or - and pileups generated with ‘--by_strand --flip_negative_strand’ and combining +- and -+ using a custom modify_2Dintervals_func in coolpup.py. Pileup heatmaps were generated using the coolpuppy function plotpup.py with settings ‘--plot_ticks’ and appropriate values for ‘--cols and ‘--rows’. ‘--norm_corners 10’ was used in all cases except Supp. Fig. 3E and Supp. Fig. 7G. Stripe plots in Fig. 3B were plotted using plotpup.py with settings ‘--stripe corner_stripe --plot_ticks --lineplot’.

### Peak and coverage files

Peaks (bed or narrowPeak) and coverage files (bigWig) used are listed in Supp. Tables 6-7. Overlaps between peaks were generated using BEDTools intersect or bioframe (Open2C et al., 2022) overlap. Peaks were randomly sampled in some cases to get the same or similar number of peaks for comparisons. Distances to the closest peak were generated using BEDTools closest and peaks were merged using BEDTools merge. For Supp. Fig. 3A-B and Fig. 2F, ENCODE DNase-seq peaks or H3K27ac peaks not overlapping TSSs, CGIs or RING1B were split into quartiles based on the signalValue column. CGIs and dinucleotide frequencies were derived from CAP-CGI data (Illingworth et al., 2010). TSSs were defined as the first TSS for all genes in refGene (O’Leary et al., 2016). Processed H3K27ac values from stimulated THP-1 derived macrophages (Reed et al., 2022) were split into quartiles based on the m0000_VST and m1440_VST columns and regions with adjusted p-values < 0.5 and going from quartiles 1 or 2 to 4 or (up-regulated) or 4 to 1 or 2 (down-regulated) were used. For Supp. Fig. 6C, compartments were called using cooltools eigs-cis at 50kb resolution and eigenvector E1 values above and below zero were called as A and B and adjacent bins in the same compartment merged. TSSs not overlapping RING1B binding sites within 50 kb of compartment edges were compared to Q4 TSSs not overlapping RING1B at least 100 kb from compartment edges. Zebrafish H3K27ac enrichment values were taken from figure 2d source data from (Yang et al., 2020) and brain/muscle enrichment defined as those with values >4 in one and <1 in the other tissue. Processed *Drosophila melanogaster* RNA-seq data from (Loubiere et al., 2020) were split into quartiles based on the baseMean value. For GM12878, the top 20000 H3K27ac peaks (based on signalValue) not overlapping H3K27me3 from ENCODE were used and overlapped with ENCODE MYC peaks (Supp. Fig. 7A) or subcompartments from (Rao et al., 2014) (Supp. Fig. 7D). GPSeq data at 100 kb resolution were downloaded from https://github.com/ggirelli/GPSeq-source-data (source data figure 2e) and values from two experiments were averaged and split into three bins, which were overlapped with CAP-CGI regions which dit not overlap H3K27me3 in HAP1 (from ENCODE). Peaks were converted between assemblies using UCSC liftOver (Hinrichs et al., 2006). Coverage heatmaps and lineplots were generated using deepTools (Ramírez et al., 2016) computeMatrix with settings ‘-reference-point -- referencePoint center -a 5000 -b 5000’ and plotted using deepTools plotHeatmap or plotProfile.

### Expression quartiles and H3K27ac

4SU-seq data from mESCs and differentiating towards NPCs (Boyle et al., 2020) were aligned to the mm10 genome in paired-end mode using STAR 2.7.1a (Dobin et al., 2013) with settings ‘--outFilterMultiMapNmax 1’. H3K27ac ChIP-seq data from mESCs (Luo et al., 2020) and NPCs (Bonev et al., 2017) were aligned to the mm10 genome in single-end mode using STAR 2.7.1a with settings ‘-- outFilterMultiMapNmax 1 --alignMatesGapMax 2000 --alignIntronMax 1 -- alignEndsType EndToEnd’. Duplicate reads were discarded using Picard (http://broadinstitute.github.io/picard/), and reads not aligning to autosomes, X, or Y were removed using samtools 1.10 (Li et al., 2009). Read counts were generated using HOMER 4.10 (Heinz et al., 2010) annotatePeaks.pl with settings ‘-noadj -len 0 -size given’ in the first exon of every refGene gene (for 4SU-seq) or merged H3K27ac peaks within 5 kb from mESCs and NPCs (for H3K27ac ChIP-seq). TMM normalisation was performed using edgeR (Robinson et al., 2010) in R 4.1.3 (R Core Team, 2022). For 4SU-seq, the mean normalised values from both replicates were averaged, divided by the length of the exon, and split into quartiles. For H3K27ac, DE analysis using limma (Ritchie et al., 2015) was performed with contrast ∼0+Sample. Peaks with adjusted p-values < 0.05 were split into those higher in mESCs or NPCs.

### DNA FISH

mESCs grown on slides were fixed in 4% paraformaledhyde, permeabilized in PBS/0.5% Triton X, dried, and then stored at −80°C prior to hybridisation. Slides were incubated in 100 μg/mL RNase A in 2× SSC for 1 h at 37°C, washed briefly in 2× SSC, passed through an alcohol series, and air-dried. Slides were incubated for 5 min at 70°C, denatured in 70% formamide/2× SSC (pH 7.5) for 40 min at 80°C, cooled in 70% ethanol for 2 min on ice, and dehydrated by immersion in 90% ethanol for 2 min and 100% ethanol for 2 min prior to air drying. 1 μg of fosmid DNA was labeled by nick translation to incorporate green-dUTP (Enzo Lifesciences), Alexa fluor 594-dUTP (Invitrogen) or digoxigenin-11-dUTP (Roche). 100 ng of each fosmid, 6 μl of Cot1 DNA per fosmid, and 5 μg of sonicated salmon sperm DNA were dried in a spin-vac and then reconstituted in 30 μl of hybridisation mix. Probes were then denatured for 5 min at 80°C and reannealed for 15 min at 37°C. Fosmid probes were hybridised to slides under a sealed coverslip overnight at 37°C. Slides were washed the next day four times for 3 min in 2× SSC at 45°C and four times for 3 min in 0.1× SSC at 60°C, and the digoxigenin labelled probe detected with anti-digoxigenin antibody (Roche) and Alexa-Fluor 647 donkey anti-sheep antibody (Invitrogen). Slides were stained with 4,6-diaminidino-2-phenylidole (DAPI) at 50 ng/mL, mounted in VectaShield (Vector Laboratories), and sealed with nail varnish. Slides were imaged on the SoRa spinning disk confocal microscope (Nikon CSU-W1 SoRa) and images were denoised and deconvolved using NIS deconvolution software (blind preset) (Nikon). 3D images are shown in the figures as maximum intensity projections prepared using ImageJ. The distances between the relevant spots was calculated using the Imaris spots function. Statistical comparison was performed using a Wilcoxon signed-rank test (scipy.stat.wilcoxon with default parameters) and multiple test correction using Benjamini-Hochberg (statsmodels.stats.multitest.multipletests with alpha=0.1). Probe coordinates are found in Supp. Table 8

---

**Supplementary figure 1.**
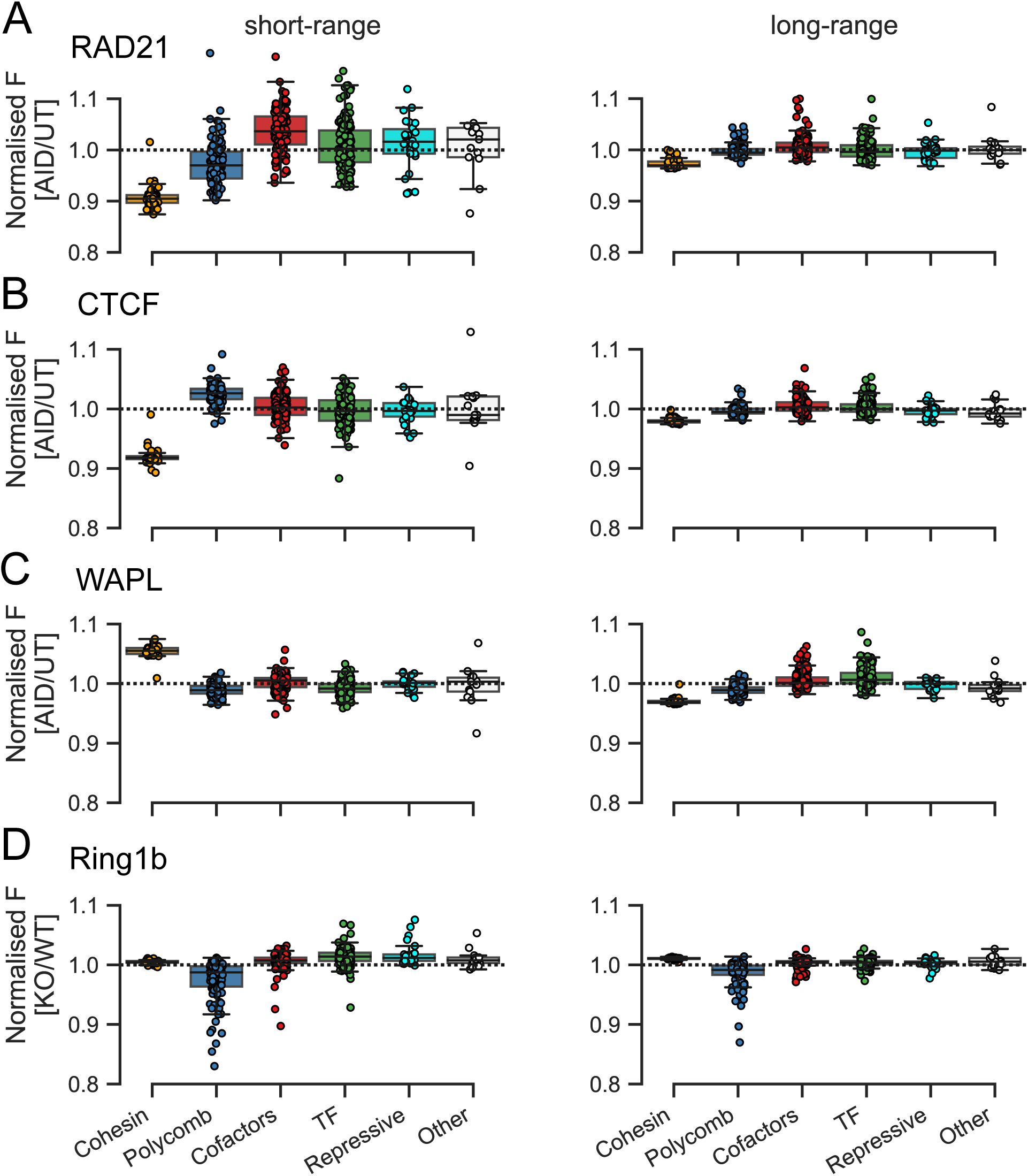
Effect of cohesin subunit degradation and Ring1b knockout on contact frequency screen enrichment values. **(A-D)** F-values for (left) short-range (0.1-1 Mb between pairs) and (right) long-range (1-10 Mb between pairs) enrichment were calculated for each Hi-C/micro-C dataset and the relative enrichment derived by dividing by the mean F for each dataset. This value was divided between treatment (AID = Auxin treated, or KO = knockout) and control (UT = Untreated, or WT) for **(A)** RAD21-AID, **(B)** CTCF-AID, **(C)** WAPL-AID, and **(D)** Ring1b KO.

**Supplementary figure 2.**
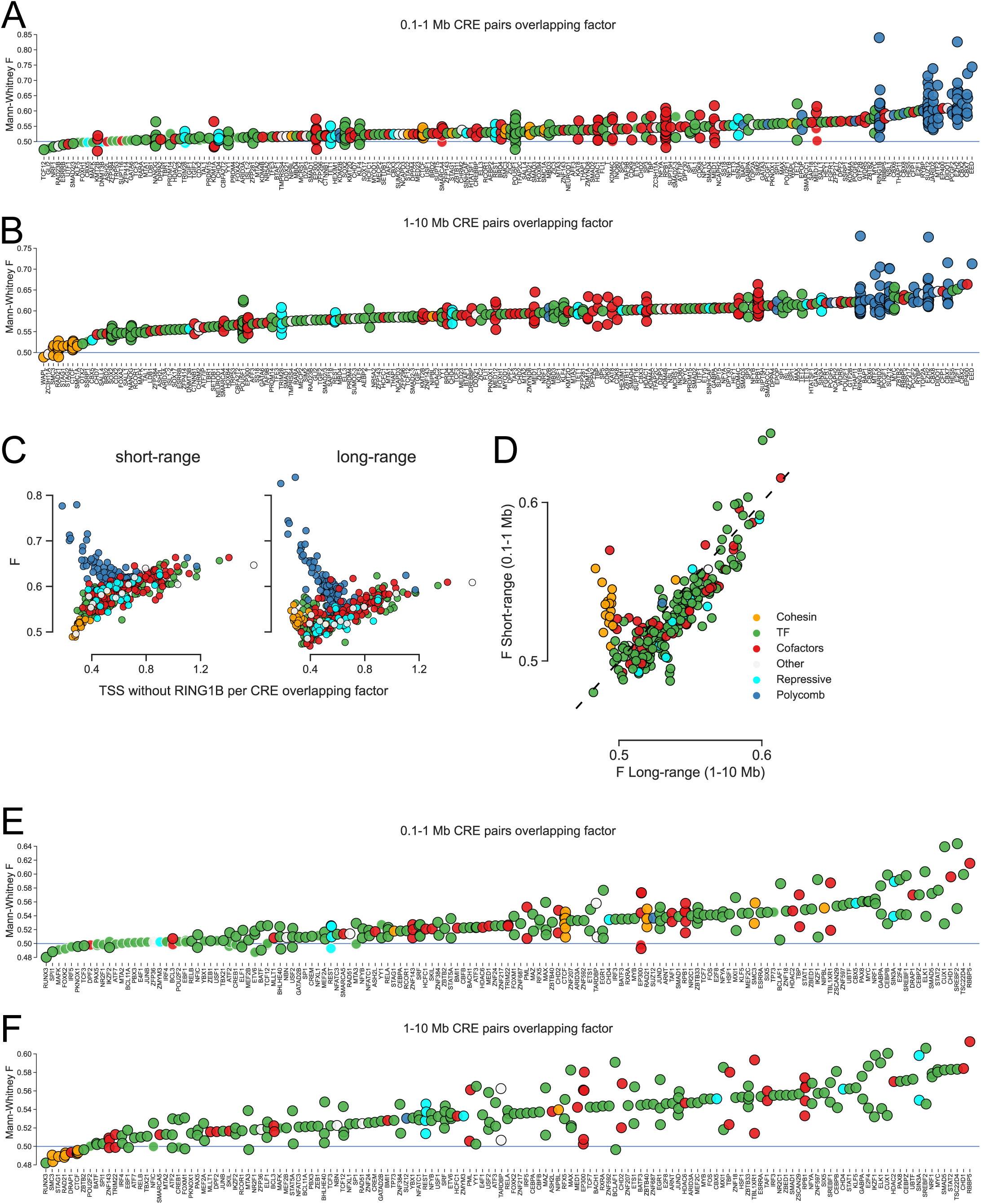
Contact frequency screen results in mESCs and human GM12878 lymphoblastoids. **(A and B)** F values for short-range **(A)** and long-range **(B)** enrichment in mESCs. Black outlined dots are significantly enriched (adjusted p<0.05), while non-outlined ones are not. **(C)** Correlation between the number of non-RING1B overlapping TSSs per CRE overlapping the factor and its enrichment (F) value at short (left) and long (right) range. **(D)** Effect sizes for factors with significantly enriched chromatin interactions compared to unbound CREs in the human GM12878 lymphoblastoid cell line. x and y axes show enrichment at short-range and long-range. Colours represent the group the factor belongs to. **(E and F)** Same as (A and B) but for GM12878.

**Supplementary figure 3.**
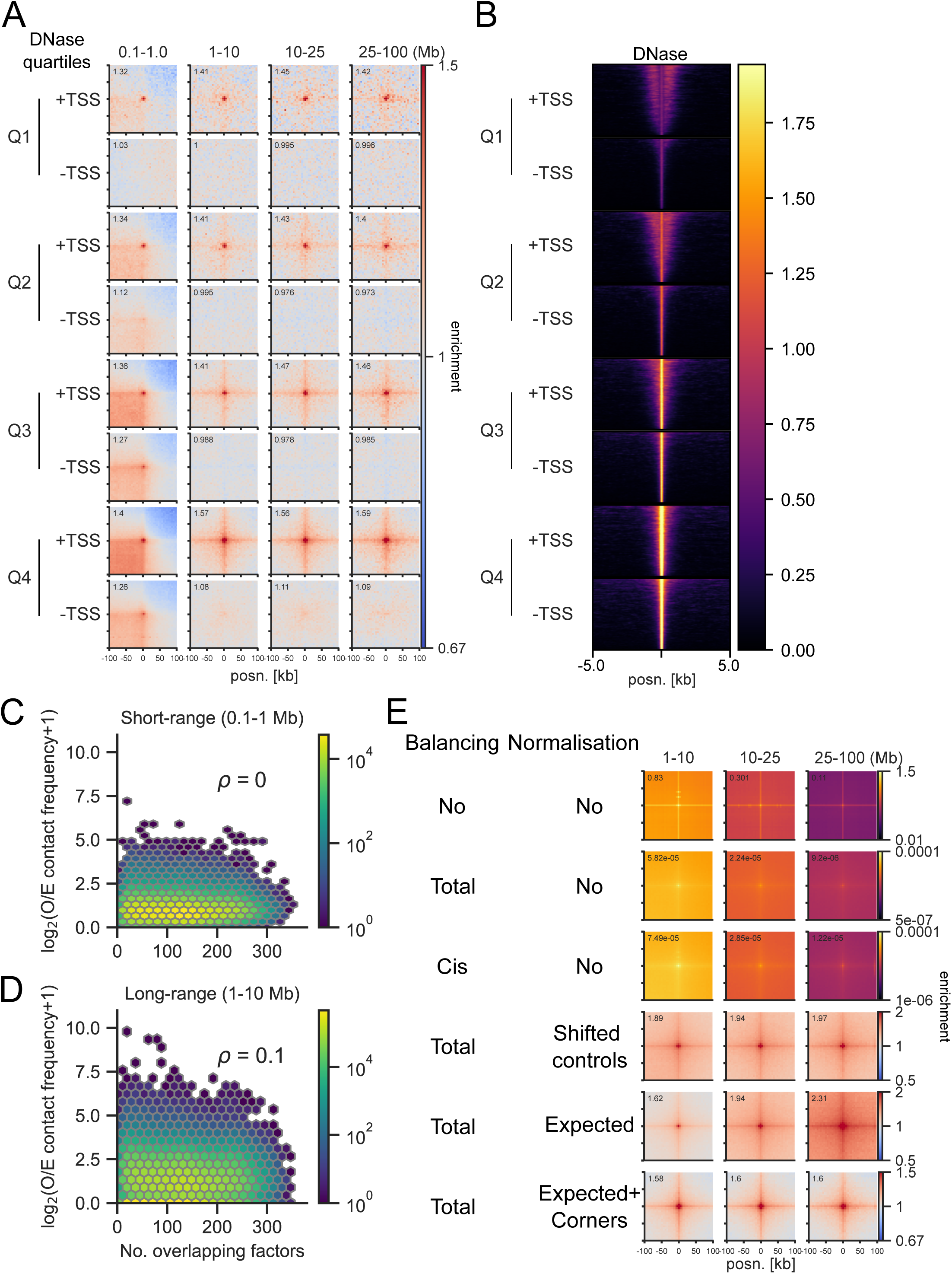
Exclusion of accessibility, binding, balancing, and normalisation as artifacts. **(A)** Pileup analysis of micro-C data from mESCs in quartiles split by DNase-seq signal and proximity to TSSs (<1KB or >5kb; 2705 peaks per group). **(B)** DNase-seq signal for regions used in (A). **(C and D)** Hexbin plots showing the correlation (ρ: Spearman’s correlation coefficient) between the sum of the total number of factors overlapping the regions on both sides and contact frequencies at short-range **(C)** and long-range **(D)**. **(E)** Pileup analysis of 5898 CFP1 peaks not overlapping RING1B in micro-C data from mESCs using different balancing and normalisation parameters. Balancing refers to ICE normalisation, based on total or only cis contacts. Normalisation is performed using either 5 randomly shifted regions for each peak or based on calculated expected values. Corner normalisation refers to normalising the total signal by the average of the signal in the 4×10 corner pixels.

**Supplementary figure 4.**
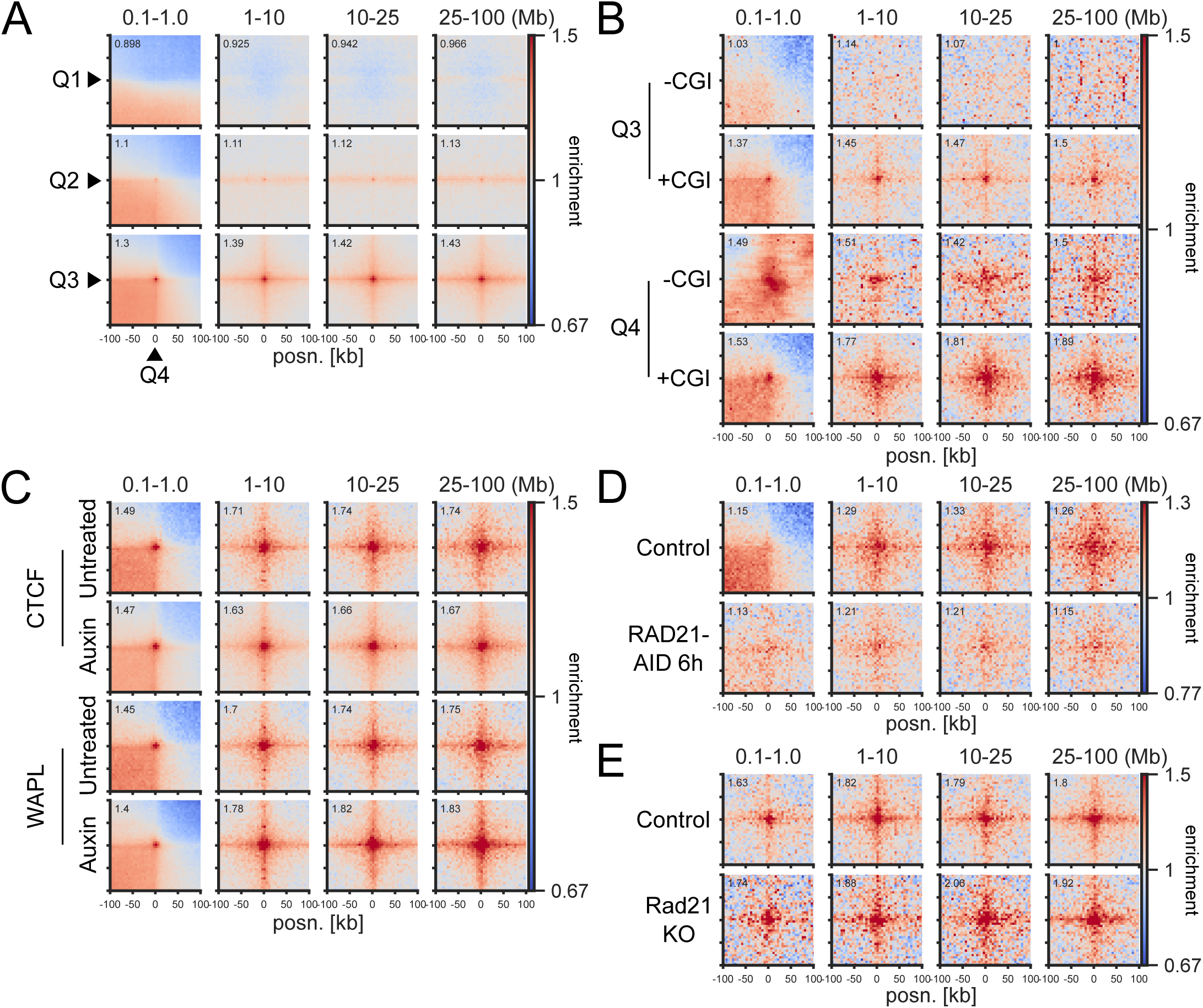
ULIs between TSSs, effect of CTCF and WAPL degradation, and effect of RAD21 long-term depletion or knockout. **(A)** Pileup analysis between different non-RING1B bound TSS quartiles in micro-C data from mESCs. **(B)** Pileup analysis of the top 2 non-RING1B bound TSS quartiles based on expression and split by overlap with CGIs (732 peaks per group) in micro-C data from mESCs. **(C)** Pileup analysis between CGI Q4 regions in micro-C data from CTCF-AID and WAPL-AID mESCs. **(D)** Pileup analysis between CGI Q4 regions in Hi-C data from RAD21-AID mESCs. **(E)** Pileup analysis between CGI Q4 regions in Hi-C data from WT and Rad21 KO thymocytes.

**Supplementary figure 5.**
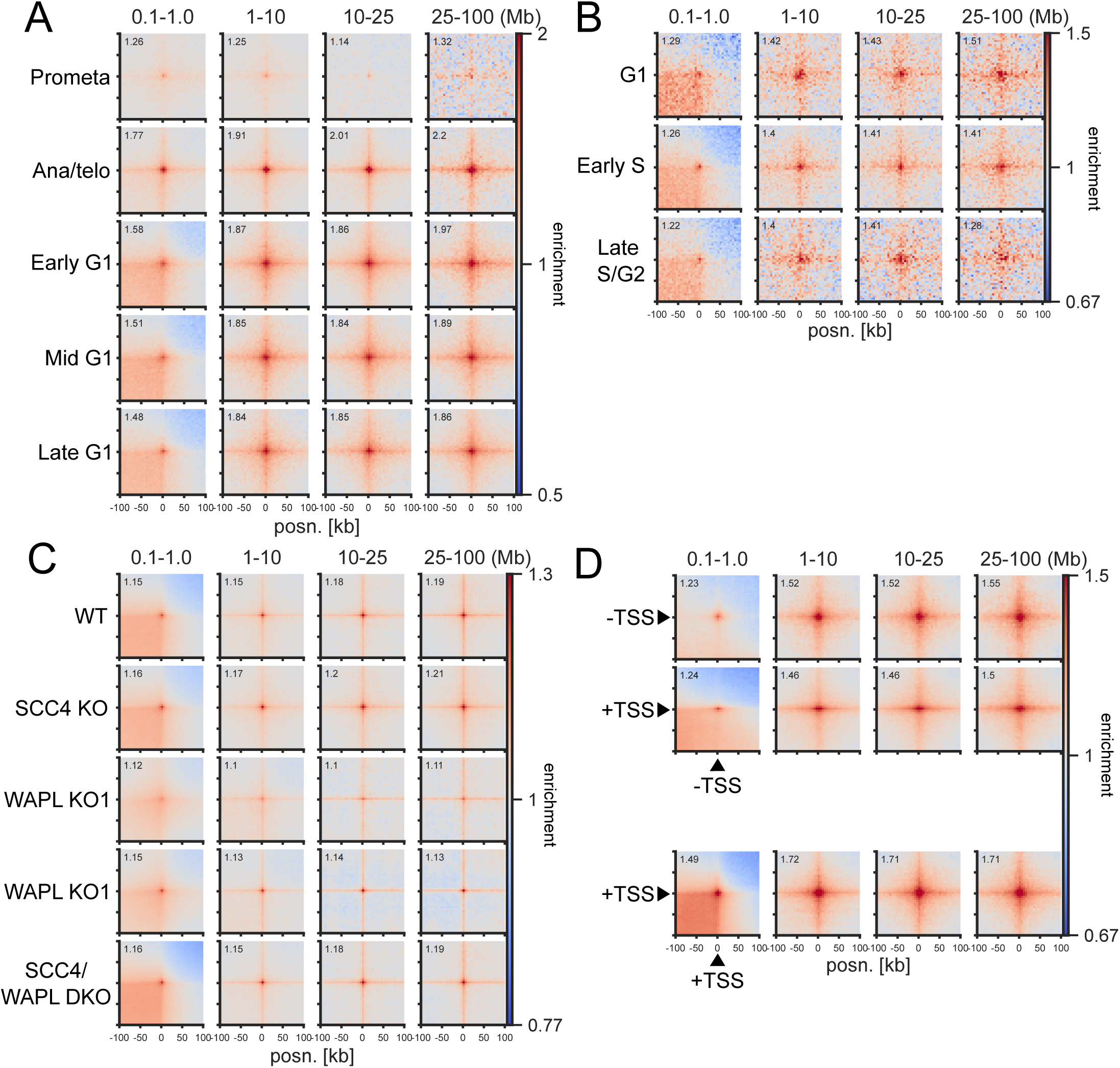
Cell cycle phase, cohesin disruption in HAP1, and TSS-non TSS interactions. **(A)** Pileup analysis between CGI Q4 regions in Hi-C data from cell cycle staged mouse erythroblasts. **(B)** Pileup analysis between CGI Q4 regions in merged single-cell Hi-C data from cell cycle stage-inferred mESCs. **(C)** Pileup analysis between high CpG density CGI (Q4) regions not overlapping H3K27me3 (22’924 Peaks) in Hi-C data from HAP1 cells with or without SCC4 and/or WAPL. **(D)** Pileup analysis between Q4 H3K27ac regions (6268 peaks; -TSS) and Q4 TSSs in micro-C data from mESCs.

**Supplementary figure 6.**
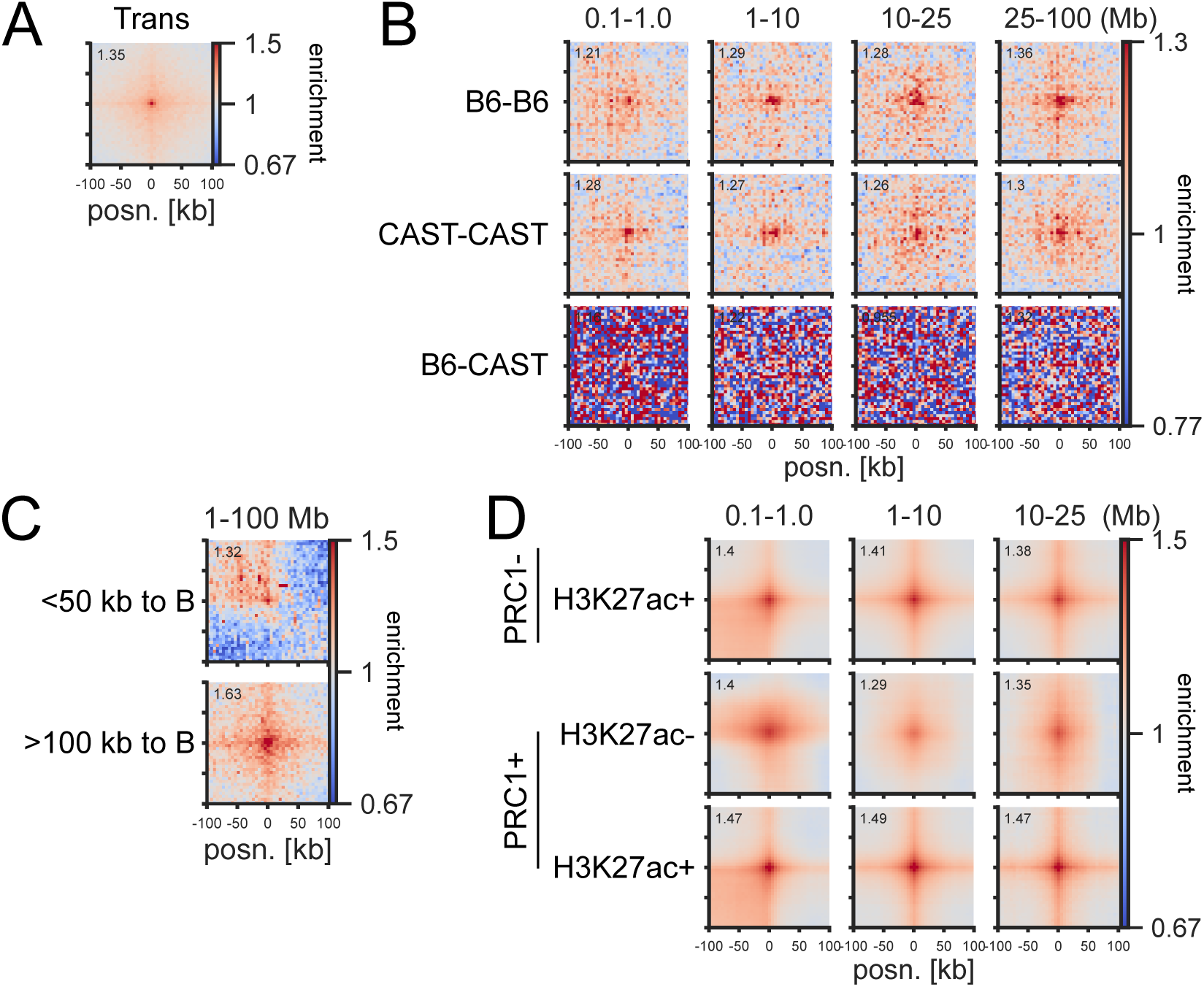
*Trans* interactions, compartment switches, and interactions in *Drosophila melanogaster*. **(A)** Pileup analysis between CGI Q4 regions on different chromosomes (*trans)* in micro-C data from mESCs. **(B)** Pileup analysis for non-RING1B bound CGI regions (11’157 peaks) in Hi-C data from SNP-phased hybrid mouse CD4 T-cells. **(C)** Pileup analysis for non-RING1B bound TSSs close to B compartments (top; 1139 peaks) or Q4 TSSs far from B compartments (bottom; 1139 peaks) in micro-C data from mESCs. **(D)** Pileup analysis for regions with different combinations of H3K27ac and PRC1 (SUZ12 and PSC) regions (4722, 3187, and 3350 peaks) in Hi-C data from *Drosophila melanogaster* eye-antennal imaginal discs.

**Supplementary figure 7.**
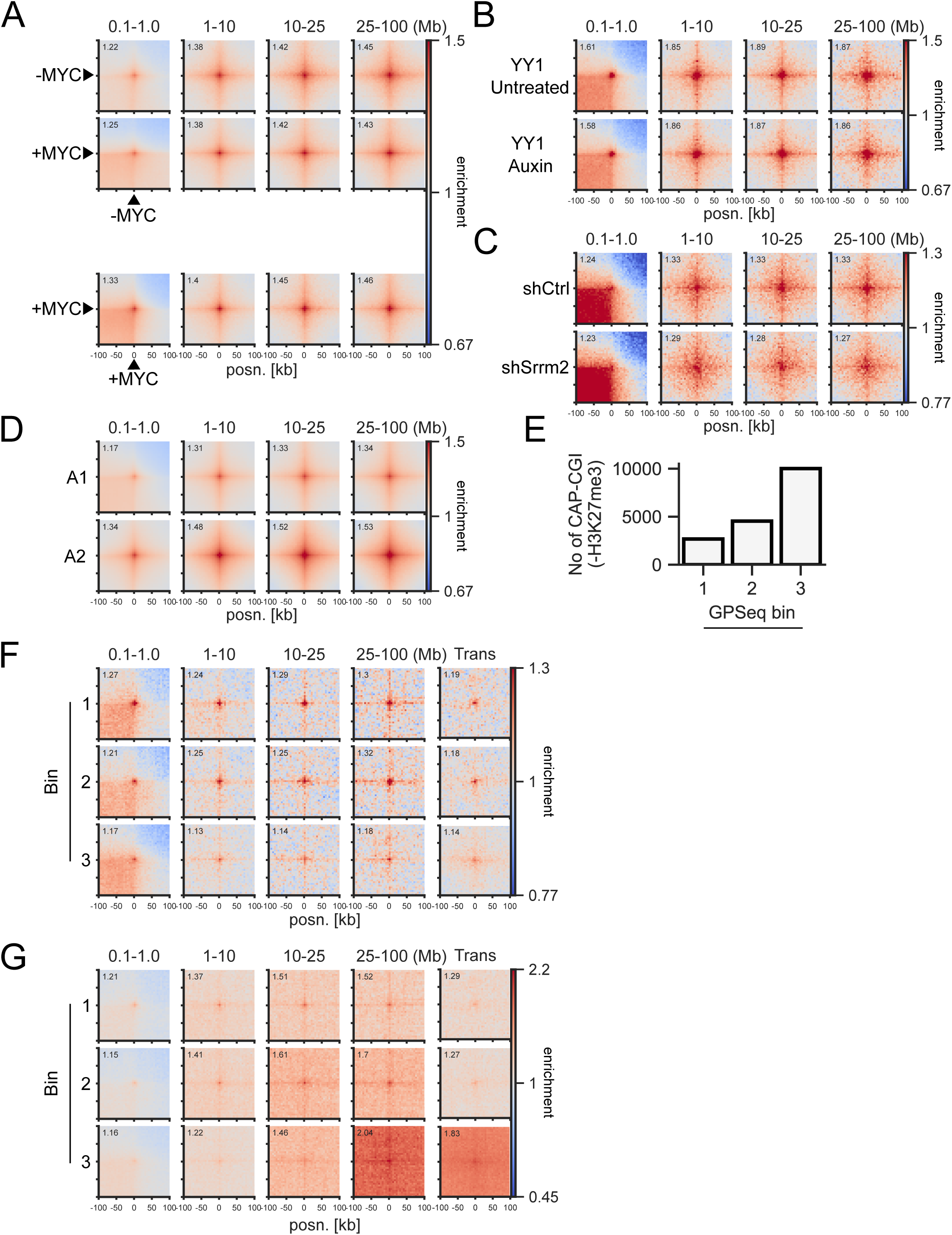
MYC binding, effect of YY1 and Srrm2 perturbation, subcompartments, and correlation with radial positioning. **(A)** Pileup analysis between H3K27ac peaks with or without MYC (3301 peaks per group) in Hi-C data from GM12878. **(B)** Pileup analysis between CGI Q4 regions in micro-C data from YY1-AID mESCs. **(C)** Pileup analysis between CGI Q4 regions in Hi-C data from AML12 cells with control or *Srrm2* shRNA. **(D)** Pileup analysis of H3K27ac peaks overlapping the A1 (9445 peaks) or A2 (8082 peaks) subcompartments in Hi-C data from GM12878. **(E)** Number of CGIs not overlapping H3K27me3 in HAP1 overlapping three bins based on GPSeq signal, where higher means more central nuclear localisation. **(F)** Pileup analysis of CGIs overlapping different GPSeq bins (2670 peaks per group) in *cis* (left) and *trans* (right) in Hi-C from HAP1 cells. **(G)** Pileup analysis of CGIs overlapping different GPSeq bins in *cis* (left) and *trans* (right) in Hi-C from HAP1 cells without corner normalisation.

## Author contributions

CRediT author statement:

E.T.F. – Conceptualisation, Data curation, Formal analysis, Funding acquisition, Methodology, Project administration, Software, Visualisation, Writing – Original Draft, I.M.F. – Conceptualisation, Data curation, Formal analysis, Methodology, Software, Writing – Review & Editing

S.B. – Investigation, Validation

W.A.B. – Conceptualisation, Funding acquisition, Project administration, Resources, Supervision, Writing – Original Draft

## Acknowledgements

E.T.F. was supported by a fellowship from the Swiss National Science Foundation (P500PB_206805). Work in the group of W.A.B. is supported by MRC University Unit grant MC_UU_00007/2. We would like to thank Hannes Becher for statistical advice. This work has made use of the resources provided by the Edinburgh Compute and Data Facility (ECDF) (http://www.ecdf.ed.ac.uk/).

